# Modeling High-grade serous ovarian carcinoma using a combination of *in vivo* electroporation and CRISPR/Cas9 mediated genome editing

**DOI:** 10.1101/2021.03.29.437221

**Authors:** Katie Teng, Mathew Ford, Keerthana Harwalkar, YuQi Li, David Farnell, Nobuko Yamanaka, Ton Nu Tuyet Nhung, David Huntsman, Jocelyne Arseneau, Yojiro Yamanaka

**Author notes:** Correspondence, Yojiro Yamanaka PhD, 1160 Pine Avenue West, Montreal, H3A1A3, Quebec, Canada.

## Abstract

Ovarian cancer remains the most lethal gynecological cancer today. High-grade serous ovarian carcinoma (HGSC) is the most common and lethal type of ovarian cancer and is most frequently diagnosed at advanced stages. Here, we developed a novel strategy to generate somatic ovarian cancer mouse models using a combination of *in vivo* electroporation and CRISPR/Cas9 mediated genome editing. We mutated tumor suppressor genes associated with HGSC in two different combinations; *Brca1, Tp53, Pten* with/without *Lkb1* and successfully generated HGSC, however, with different latencies and pathophysiology. By utilizing Cre lineage tracing in our system, we visualized peritoneal micrometastases in an immune-competent environment. Because our strategy is flexible in selecting mutation combinations and targeting areas, it would be useful for generating ovarian cancer mouse models.

**Significance Statement:** High grade serous ovarian cancer (HGSC) is the most common ovarian malignancy but our knowledge of early tumorigenesis is still quite limited due to late diagnosis of patients. We developed a new strategy of generating mouse ovarian cancer models using a combination of in vivo electroporation and CRISPR-mediated genome editing. We demonstrated that a combination of three tumor suppressor gene mutations (*Brca1, Tp53 and Pten*) is sufficient to develop HGSCs. Interesting, an additional mutation in *Lkb1* drastically changed tumor latency, penetrance and pathophysiology through changing its cell-of-origin. Our strategy is highly flexible in selection of mutation combinations and targeting areas in immune competent mice and useful to study early tumorigenesis of ovarian cancer.

## Introduction

According to the American Cancer Society, 21,750 women in the United States will be newly diagnosed with Ovarian Cancer (OC) in 2020 and 13,940 will succumb to this devastating disease^1^. It is ranked fifth in cancer deaths among women and is considered the most lethal gynecological disease^1^. High-grade serous ovarian carcinoma (HGSC) is the most common type of OC and is the cause of ~70-80% of all OC deaths^2,3^. Due to the asymptomatic nature of this cancer, the majority of these patients are diagnosed at advanced stages (III or IV) after metastasis to the peritoneal cavity has occurred^3,4^.

Despite its name, the origin for HGSC remains controversial^5,6^. The ovarian surface epithelium (OSE) has been the presumed cell-of-origin for HGSC for many years^8^. However, over recent years, the notion that the cell-of-origin for HGSC resides in the OSE was challenged by the absence of a clearly defined precursor lesion in both the ovary and OSE in patients^9^. High-risk patients, such as *BRCA1* and *BRCA2* mutation carriers, often undergo prophylactic salpingo-oophorectomies in which both their fallopian tubes (FTs) and ovaries are removed^10,11^. Interestingly, by extensively examining the distal end of these FTs using the “sectioning and extensively examining the fimbriated end” (SEE-FIM) protocol, microscopic precancerous lesions such as serous tubal intraepithelial carcinomas (STICs) were identified^12–14^. Genomic studies have revealed that these precursor lesions carry the same mutations as the HGSC tumours that form. These data suggests the distal FTE can be a cell-of-origin for HGSC^12,15,16^. However, in sporadic cases of HGSC, STICs are only identified in 50-60% of patients while the remaining 40%-50% are undetectable^9^. It is unknown whether this is simply due to inaccurate detection or if there is another source for HGSC tumorigenesis^17,18^.

HGSC tumours display significant genomic heterogeneity and are driven by copy number alterations as few somatic mutations are detected in the tumours^9,19^. The most common recurrent mutation detected are *TP53* mutations found in 96% of the tumours while mutations in *BRCA* genes and other homologous recombination repair genes account for a total of 50% of HGSC cases^9,19^. Additionally, several distinct copy number signatures have been identified that are linked to molecular pathways and drug responsiveness^20,21^, suggesting that the multi-mutational processes in single HGSC patients contribute to the complex evolution of HGSC subtypes. Although recurrent amplification and deletion of chromosomal loci have been reported in several papers ^21–23,24^ the genes and pathways driving cell proliferation and metastasis have not been fully explored. Alterations of single chromosomal loci affect many genes and currently there has not been an effective strategy to identify the genes important for driving tumorigenesis and pathophysiology of HGSCs.

Developing genetically engineered mouse models (GEMMs) have proven to be effective in understanding cancer initiation and progression^25^. These models allow for the study of gene function *in vivo* and elucidate the pathways involved in early tumourigenesis^25^. However, traditional germline GEMMs are limited by availability of lineage specific Cre mouse lines and conditional alleles of the gene-of-interest. In addition, generation of desired genetic combinations in germline GEMMs is costly and can be time-consuming. In ovarian cancer modeling, *Pax8-TetOn-Cre* and *Ovgp1-CreERT2* systems were used successfully to generate HGSCs from the FTE, however, these systems would not be suitable to examine various allelic combinations and screening for other candidate genes involved in HGSC tumorigenesis.

Here we present an *in vivo* fallopian tube electroporation method that can be used to target the distal murine fallopian tube with high flexibility in selecting gene targets. CRISPR/Cas9 and Cre plasmids were directly injected into the fallopian tube lumen and were delivered to the fallopian tube epithelial cells via electroporation. By using this method, we successfully generated a model for HGSC by targeting *Brca1, Tp53* and *Pten* and showed that with the addition of *Lkb1* loss, the cell of origin can vary and significantly decrease tumor latency while increasing penetrance. This indicates the cell type specific susceptibilities to malignant transformation and the subsequent link between cell-of-origin, combination of gene mutations and pathophysiology of HGSCs. Additionally, by combining Cre mediated lineage tracing, we were able to visualize peritoneal micrometastasis which can be advantageous in studying the complex interaction between host and tumor cells during peritoneal implantation of HGSC. The system developed has the potential to be a flexible and powerful tool for understanding malignances arising from the female reproductive system in an immune competent environment.

## Results

### *In vivo* fallopian tube electroporation successfully delivered Cre and CRISPR plasmids into distal fallopian tube luminal epithelial cells for efficient genome manipulation

Development of the CRISPR/Cas9 system has permitted direct *in vivo* genome editing of somatic cells to generate GEMMs of cancer^24^. Several somatic GEMMs of various types of cancers have been reported using different delivery methods, such as lenti- and adeno-viral mediated delivery for the lung, hydrodynamic delivery to the liver and electroporation for the pancreas and developing brain^24^. The advantage of these approaches is the minimum needed requirement of specific mouse lines and flexibility in selecting gene mutation combinations and targeting locations.

To generate somatic GEMMs of ovarian cancer, we developed an *in vivo* fallopian tube electroporation method to deliver multiple DNA plasmids to the epithelium of the distal end of the fallopian tube (Fig 1A). To test its delivery efficiency, a solution of Cre plasmid was directly injected into the lumen of the distal fallopian tube of 4-6 week old *Rosa-LSLtdTomato* female mice. As a consequence of Cre excision, tdTomato (hereafter referred to as RFP) positive cells were observed in the distal region of the fallopian tube (Fig 1B,E). The RFP+ cells were distributed stochastically throughout the luminal epithelium and targeted both PAX8+ secretory and acetylated tubulin (AT)+ multiciliated cells, but not the stromal tissue compartment (Fig 1C-D).

**Figure 1:**
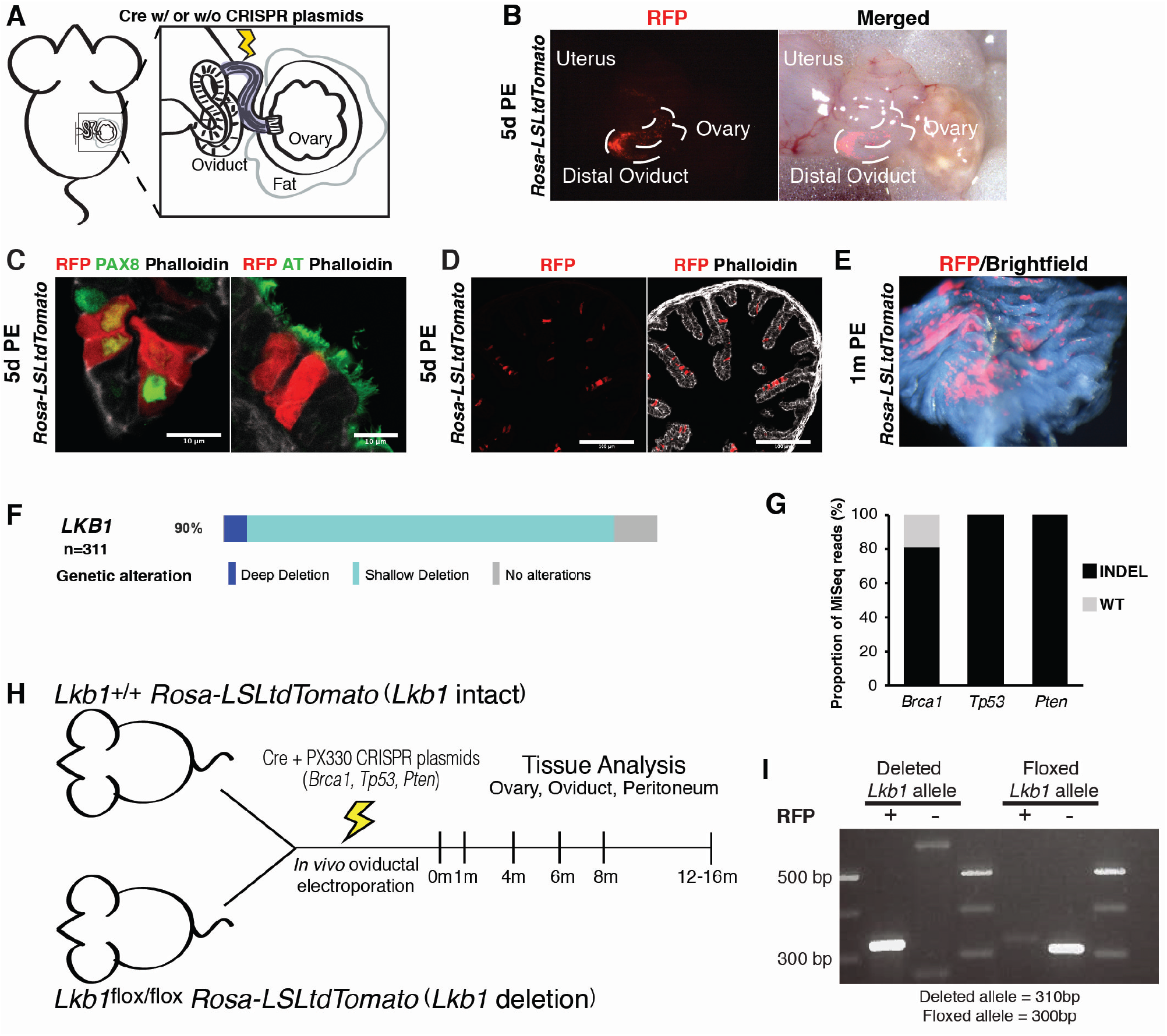
*In vivo* fallopian tube electroporation induced Cre-mediated excision in distal fallopian tube epithelial cells of *Rosa-LSLtdTomato* mice. (A) A diagram of the *in vivo* fallopian tube electroportation procedure. Fallopian tube/ovary was surgically exposed using a 1cm dorsal incision. A DNA plasmid solution was injected into the fallopian tube lumen and electroporated into fallopian tube epithelial cells. (B) RFP+ cells within the distal fallopian tube epithelium 5 days PE in *Rosa*-*LSLtdTomato* mice. (C) Transverse sections of *Rosa*-*LSLtdTomato* distal fallopian tube 5 days PE. Left panel: PAX8 (green) and phalloidin (white). Secretory cells marked by PAX8 were RFP+. Right panel: AT (green) and phalloidin (white). Multiciliated cells marked by AT were RFP+. Scale bar 10 μm (D) Transverse section of *Rosa*-*LSLtdTomato* distal oviduct 5 days PE stained with phalloidin (white). RFP+ cells were found within the luminal epithelium. Scale bars 100 μm. (E) Butterfly dissection of Cre transfected regions in distal oviduct 1m PE in *Rosa*-*LSLtdTomato* mouse. (F) Oncoprint of *LKB1* alterations in 311 patients with ovarian serous cystadenocarcinoma. 90% of these patients showed either a shallow or deep deletion in *LKB1*. (G) Frequency of indel mutations in MiSeq reads in RFP+ fallopian tube epithelial cells. (H) Schematic representation of electroporation and tissue analysis experiments in *Lkb1* intact and *Lkb1* deletion cohorts. (I) Cre-induced Lkb1 deletion allele was only detected in FACS sorted RFP+ fallopian tube epithelial cells but not in RFP-cells 1m PE.

To evaluate the effectiveness of our approach for HGSC modeling, we targeted three tumor suppressor genes, *Brca1, Tp53* and *Pten*, previously used to generate germline GEMMs of HGSC using the *Pax8-TetOn-Cre* system^8^. In addition to these three tumour suppressor genes, we selected an additional tumor suppressor gene, *LKB1*(also known as *STK11*). *LKB1/STK11* is an evolutionarily conserved pleiotropic kinase that regulates cell polarity, cell cycle and energy metabolism and is deleted in many cancers^26,27^. Ch19p13.3, where the *LKB1* locus resides, is identified as one of many recurrent chromosomal deletions in HGSC^19,21^. Somatic mutations in *LKB1* are not frequent, however, 90% of HGSCs are significantly associated with either a shallow (monoallelic loss) or deep deletion in *LKB1* (Fig 1F). Additionally, downregulation of LKB1 protein is shown to be characteristic of HGSC tumours, suggesting that loss-of-LKB1 is involved in HGSC initiation and progression^4^. Furthermore, it has been shown that loss-of-*Lkb1* impairs epithelial integrity and causes spontaneous cellular extrusions from epithelium^28^. This process of cellular extrusion and anoikis resistance is compelling in the context of early/late cancer cell dissemination in HGSC.

PX330 plasmids encoding sgRNAs against *Brca1, Tp53* and *Pten* were electroporated along with Cre plasmid into the distal fallopian tube luminal epithelium of 4-6 week old females; *Rosa-LSL-tdTomato* and *Lkb1*^flox/flox^; *Rosa-LSL-tdTomato*, which will be referred to as *Lkb1* intact and *Lkb1* deletion cohorts, respectively, hereafter (Fig 1H). To determine if indel mutations were efficiently introduced into the electroporated cells, we isolated RFP+ and RFP-fallopian tube epithelial cells 1 month PE. The targeted sequences were PCR amplified and analyzed using Sanger and MiSeq sequencing. We found that only RFP+ cells had CRISPR induced indel mutations in *Brca1, Tp53* and *Pten* while no mutations were detected in the RFP-cells. In the isolated RFP+ epithelial population, 100% of the sequence reads were indel mutations in *Tp53* and *Pten* whereas only 81% were indel mutations in *Brca1* (Fig 1G). Similarly, Cre-mediated deletion in *Lkb1* was only detected in the RFP+ population but not in the RFP-population (Fig 1I). This suggested that multiple plasmids were delivered into single electroporated cells to efficiently introduce CRISPR-mediated indel mutations and Cre-mediated *Lkb1* deletion and RFP activation.

### *Lkb1* intact and *Lkb1* deletion cohorts both developed HGSC with different latencies and penetrance

Following *in vivo* electroporation, we examined the distal fallopian tube and ovary of asymptomatic mice 4 months PE under a fluorescent dissecting scope. In both *Lkb1* intact and *Lkb1* deletion cohorts, RFP+ cells were observed in the distal fallopian tube as expected (Fig 2A). Interestingly, in only the *Lkb1* deletion cohort, papillary tumours were developed on the ovarian surface (Fig 2A,B). The cells in these tumours were highly proliferative as indicated by Ki67 staining (Fig 2C). Within 6 months PE, widespread peritoneal metastasis formed in the *Lkb1* deletion cohort and some mice generated abdominal ascites by 7 months PE which is a phenotype seen in approximately 30% of human HGSC patients^29^ (Fig 2D, Table 1). Between 6-14m PE, the incidence of peritoneal metastasis was 96% and the incidence of abdominal ascites was 74% in the *Lkb1* deletion cohort (Table 1). On the other hand, mice in the *Lkb1* intact cohort developed HGSC at a significantly lower penetrance (21%) and later onset (Table 1). Ovarian tumours were observed 9m PE and peritoneal metastasis at 11-16m PE (Table 1). In both cohorts, peritoneal metastatic tumours were generally widespread however, larger tumours were often found in the omentum and mesentery, similar to human patients^30,31^ (Fig 2E). Interestingly, two patterns of peritoneal metastasis were observed; ‘miliary’ (9/20 cases) and ‘oligometastatic’ (11/20 cases) and recapitulated patterns seen in human patients (Fig 2F)^32,33^. The ‘miliary’ pattern showed numerous millet-like lesions spreading over a wide surface of the peritoneum. This pattern is a strong negative factor with respect to overall survival in human patients. In contrast, the ‘oligometastatic’ pattern exhibited several larger tumor nodules. These two types of metastases were not mutually exclusive and both were present in single mice similar to human patients.

**Figure 2:**
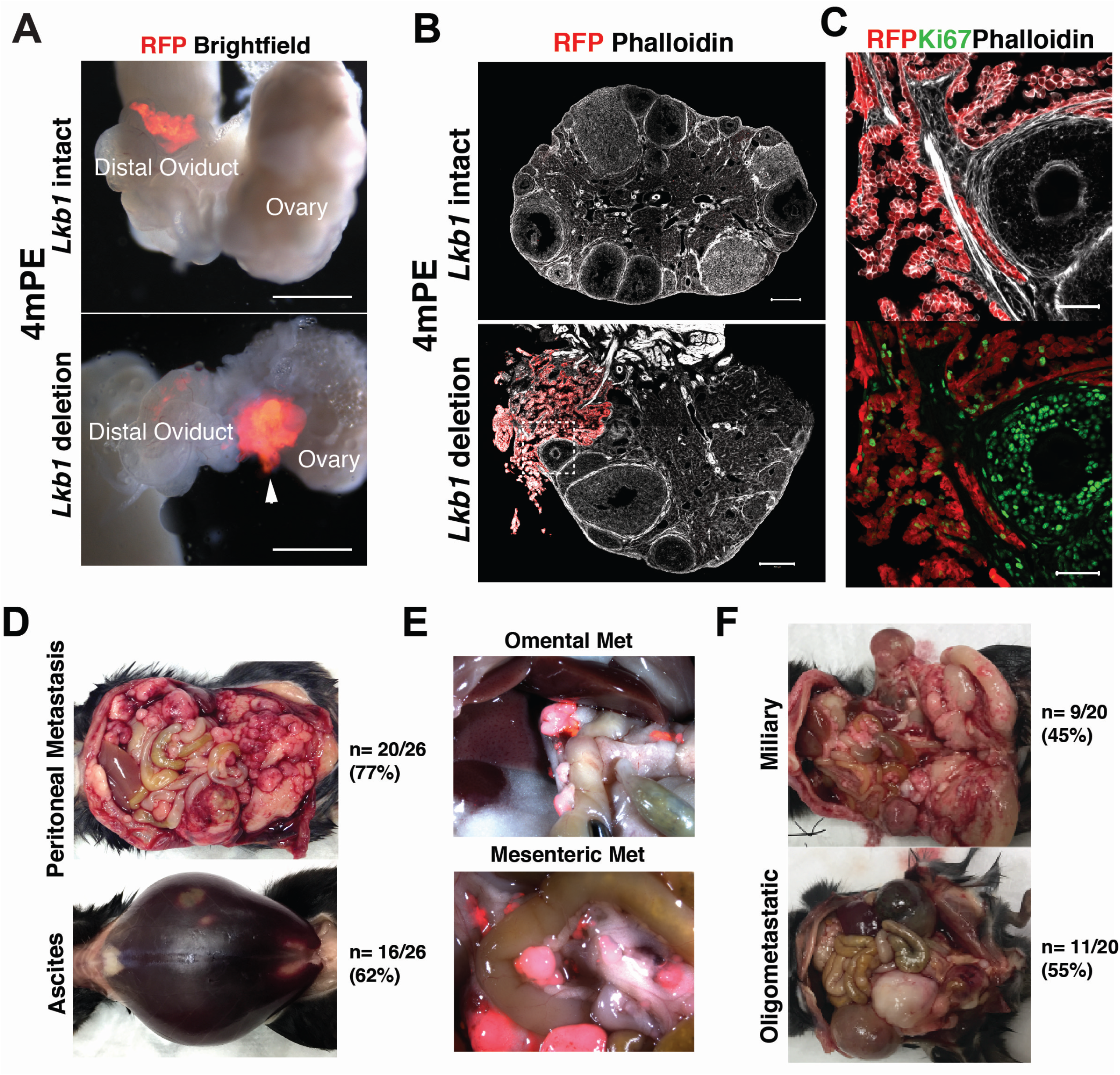
Formation of papillary ovarian tumours, peritoneal metastasis and ascites in *Lkb1* deletion cohort. (A) Dissection scope images of ovary and fallopian tube of *Lkb1* intact and *Lkb1* deletion mice 4m PE. RFP+ ovarian tumour (white arrowhead) was only detected on the ovary of an *Lkb1* deletion female. Scale bars 500 μm. (B) Sections of ovaries from *Lkb1* intact and *Lkb1* deletion mice 4m PE stained with phalloidin (white). Papillary tumour (red) on the ovary of a *Lkb1* deletion female. Scale bars 200 μm. (C) Higher magnification image of inlet marked in B. Many cells in the papillary tumour were Ki67+ (green). Scale bars 50 μm. (D) Peritoneal metastasis and abdominal ascites formation. (E) RFP+ metastatic omental and mesenteric tumours. (F) Two types of peritoneal metastasis (military and oligometastatic) in *Lkb1* deletion mice.

**Table 1.**
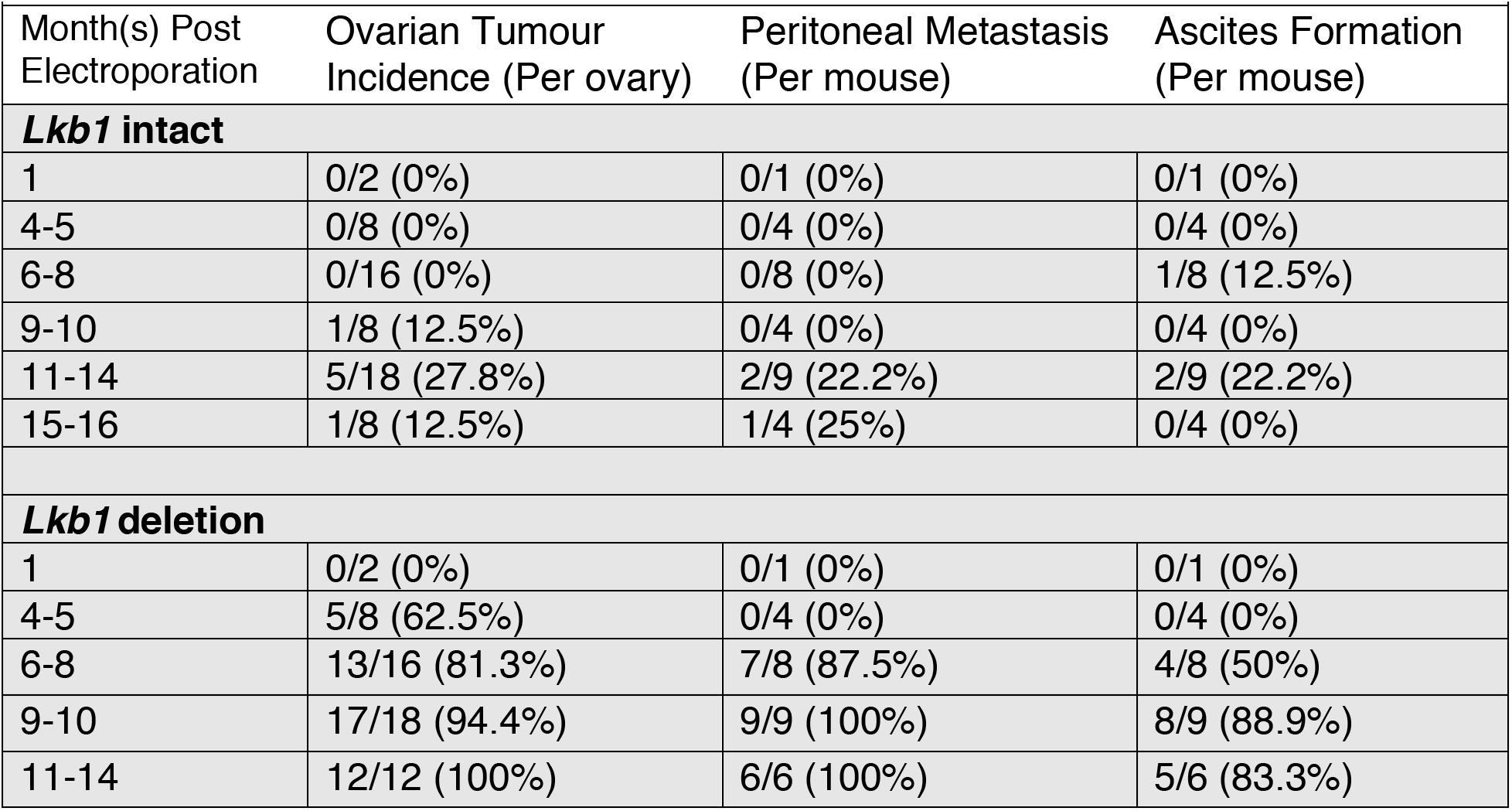
Summary of disease progression in *Lkb1* intact and *Lkb1* deletion cohorts.

Peritoneal micrometastases are undetectable by conventional imaging analysis such as MRI, PET or CT scans and via eye-inspection during surgery due to their small size. Taking advantage of our lineage tracing strategy we investigated the behavior of metastatic cells in the peritoneal environment. In a *Lkb1* deletion female 6 months PE, a relatively small ovarian tumor was formed but there were no peritoneal metastatic nodules visible to the eye or ascites formation (Fig 3A). However, under a fluorescent dissecting microscope, we detected numerous RFP+ micrometastases in the peritoneum (Fig 3A). The surface view of a relatively large RFP spot on the mesentery above the fat tissue revealed papillary tumour development on the surface of the peritoneum (Fig 3B). RFP+ cells formed a sheet-like structure with a rough surface and a papillary tumour protruded out into the peritoneal cavity. These papillary tumors were fragile and easily broken down to floating multicellular aggregates (Fig 3C). We also identified small clusters of RFP+ cells on the surface of the peritoneum, consisting of a few to a hundred RFP+ cells (Fig 3D-F). These RFP+ clusters showed a packed morphology with a clear smooth peripheral edge in contrast to jagged cell-cell contact between mesothelial cells. The disc-like clusters often had an indentation at the center and appeared to penetrate the basal membrane underneath the mesothelium (Fig 3D,E,G). Recruitment and activation of innate immune cells (e.g. LYVE1+ tissue residential macrophages) was observed immediately underneath the clusters (Fig 3D-F). Stress fiber like F-actin structure at the bottom of the RFP+ cells would suggest activation of integrins and ECM remodeling (Fig 3F).

**Figure 3:**
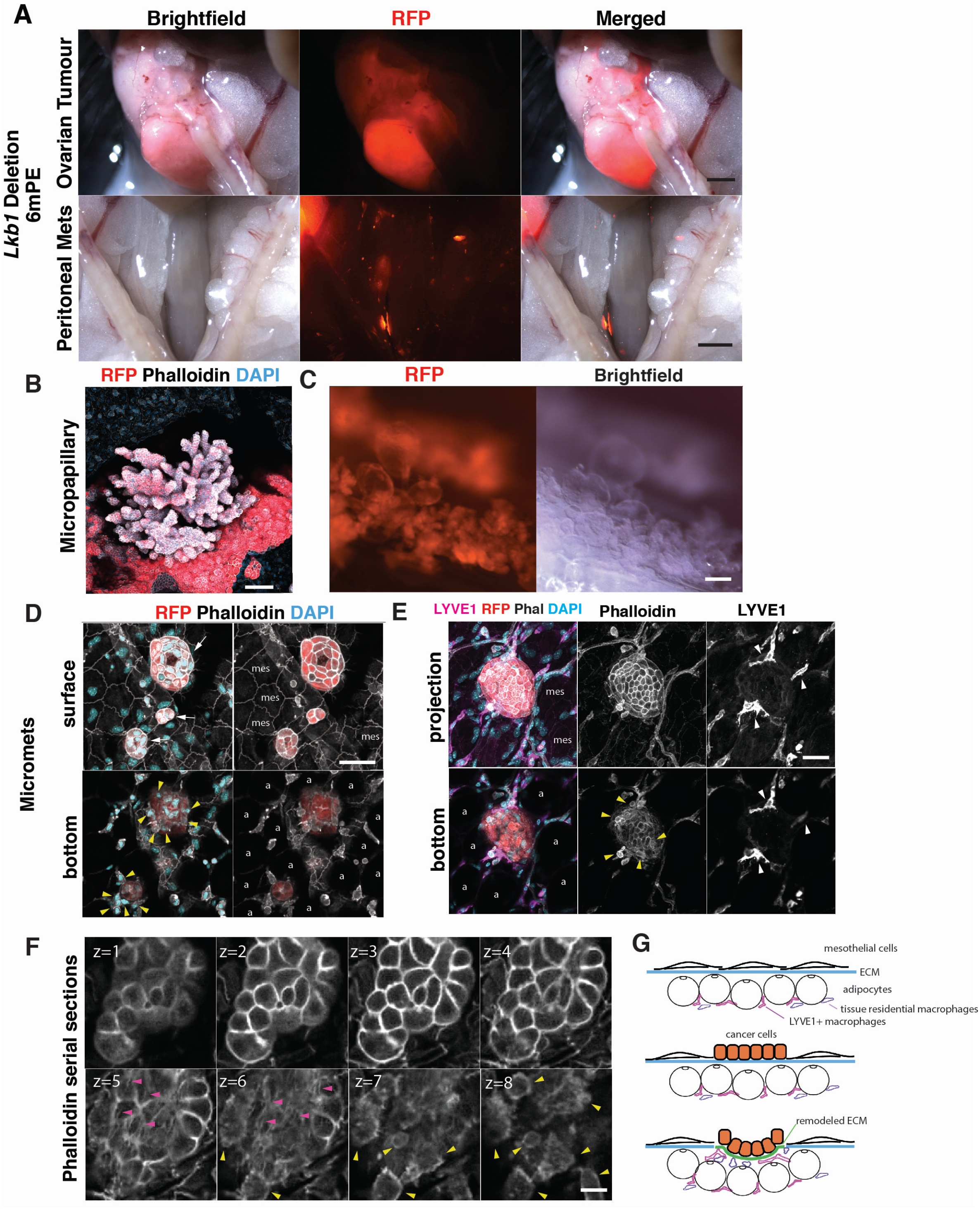
Visualization of peritoneal micrometastases. (A) Dissection scope images of primary ovarian tumour and peritoneal micrometastases in an *Lkb1* deletion female 6m PE. Scale bars 5 mm. No visible metastatic nodule in the brightfield image while numerous micrometastases were detected in the RFP image of the peritoneum. (B) A confocal projected image of a micropapillary metastasis on the peritoneum. Scale bar 100 μm. (C)Papillary micrometastases spread on the peritoneum. These tumours were fragile and easily broken down to floating multicellular aggregates. Scale bar 100 μm. (D) Three micrometastases on the peritoneum. Surface and bottom section images. Surface mesothelial cells (MES) show characteristic jagged cell-cell contact and flat cell morphology. The micrometastases (white arrows) formed a packed disc-like morphology with a central indentation. Underneath the metastases, macrophage-like cells were observed (yellow arrowheads). Adipocytes (a). Scale bar 50 μm. (E) LYVE1+ (pink) tissue residential macrophages near micrometastases (white arrowheads). Protrusive structures from LYVE1+ cells spread under the disc-like micrometastasis. Scale bar 50 μm. (F) Phalloidin serial confocal section images. 1.5 μm section intervals. Stress-fiber like F-actin structure at the bottom of the micrometastasis (Z=5,6. magenta arrowheads). Scale bar 10 μm (G) A diagram of peritoneal metastasis formation. (top) Healthy peritoneum. Thin mesothelial cells on the ECM membrane (blue) lining the visceral peritoneum in omentum and mesentery. (middle) Mesothelial clearance. Cancer cells integrate the mesothelial layer. (bottom) Tissue residential macrophages (LYVE1+) are recruited. The cancer cells and the recruited macrophages remodel the ECM underneath the cancer cells (green).

Immunohistochemical analysis and pathological review were performed on ovarian and peritoneal tumours in both cohorts. In the *Lkb1* intact cohort, these tumours were papillary and expressed PAX8, WT1 and CK8 (Fig 4A), similar to HGSC profiles in human patients and consistent with the previously reported animal models^8,34^. In the *Lkb1* deletion cohort, most tumours also showed papillary architecture. 15% of these tumours expressed PAX8, WT1 and CK8 (Fig 4B) however a significant portion (80%) of them were PAX8-, WT1+ and CK8+ (Fig 4C). It was also noted that 26% of these mice developed a rare form of ovarian cancer called ovarian carcinosarcomas or malignant mixed Müllerian tumours (MMMTs) (Fig. 4D) which was also reported in the *Ovgp1-iCreER^T2^* model by Zhai *et al*^34^. Here we have shown that HGSC tumorigenesis can be induced by targeting *Brca1*, *Tp53* and *Pten* in the oviductal epithelial cells using *in vivo* electroporation. Interestingly, loss-of-*Lkb1* along with mutations in *Brca1*, *Tp53* and *Pten* facilitated HGSC initiation and progression and these mice often presented with abdominal ascites however, many of these tumours were PAX8-.

**Figure 4:**
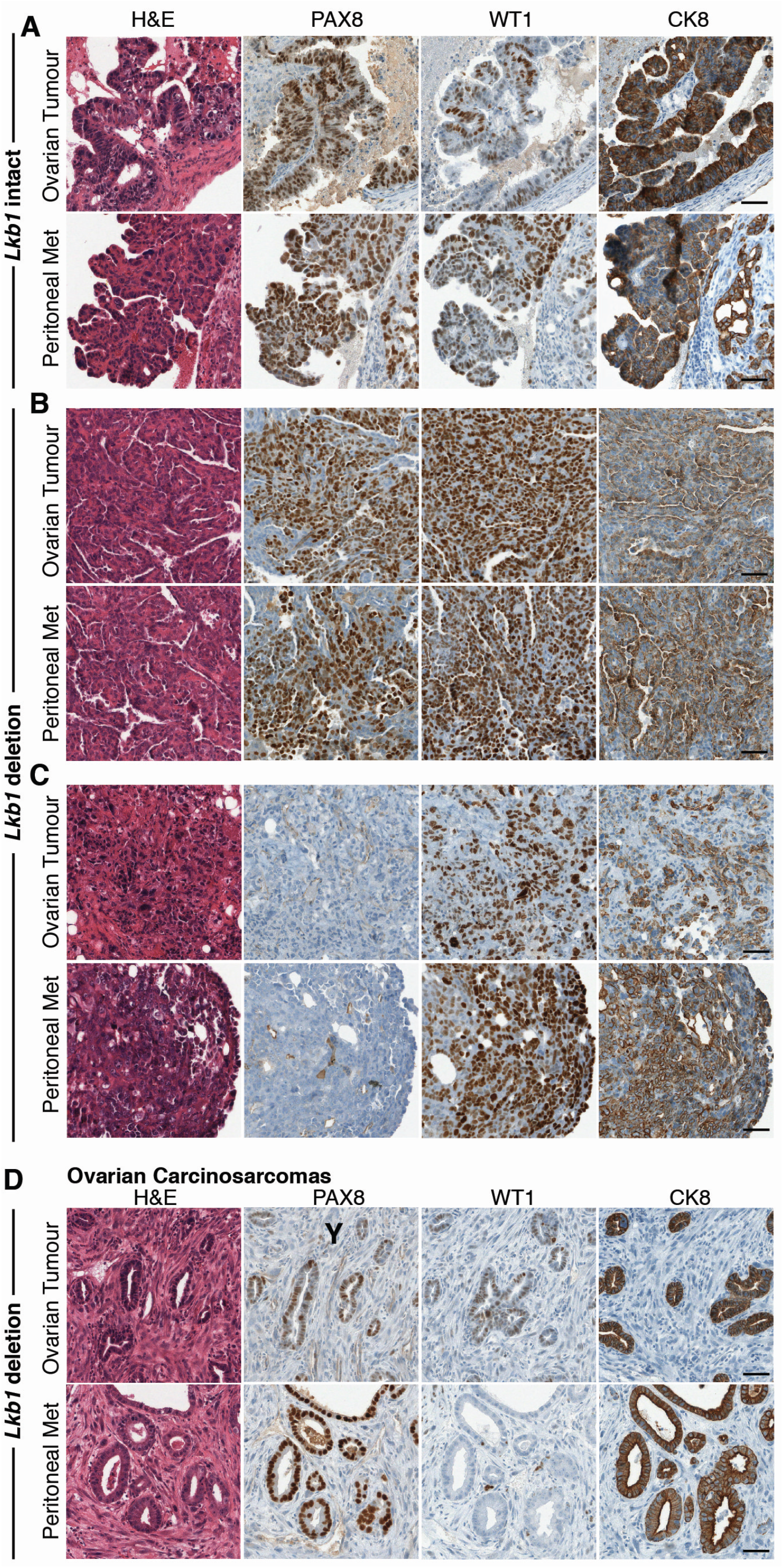
Histology of HGSC tumours and metastatic peritoneal tumours in *Lkb1* intact and *Lkb1* deletion cohorts. H&E, PAX8, WT1 and CK8 immunohistochemical staining in ovarian and peritoneal tumours. (A) *Lkb1* intact mice. Ovarian and peritoneal tumours expressed PAX8, WT1 and CK8. (B) *Lkb1* deletion mice. Ovarian and peritoneal tumours expressed PAX8, WT1 and CK8 (C) *Lkb1* deletion mice. Ovarian and peritoneal tumours expressed WT1 and CK8 but not PAX8. (D) Ovarian carcinosarcomas in *Lkb1* deletion mice. Scale bars 50 μm.

### Frameshift mutations of the targeted tumor suppressor genes were predominantly selected in ovarian tumours and peritoneal metastases

CRISPR-induced indel mutations are relatively random^25,35,36^, therefore, the genotype of individual electroporated cells soon after electroporation can, theoretically, vary in our models. If a combination of mutations in a cell gains a proliferative and/or survival advantage, the cell can form a clone in the epithelium and with further genetic and epigenetic alterations, some clones will progress to form ovarian tumours and peritoneal metastasis (Fig 5A).

**Figure 5:**
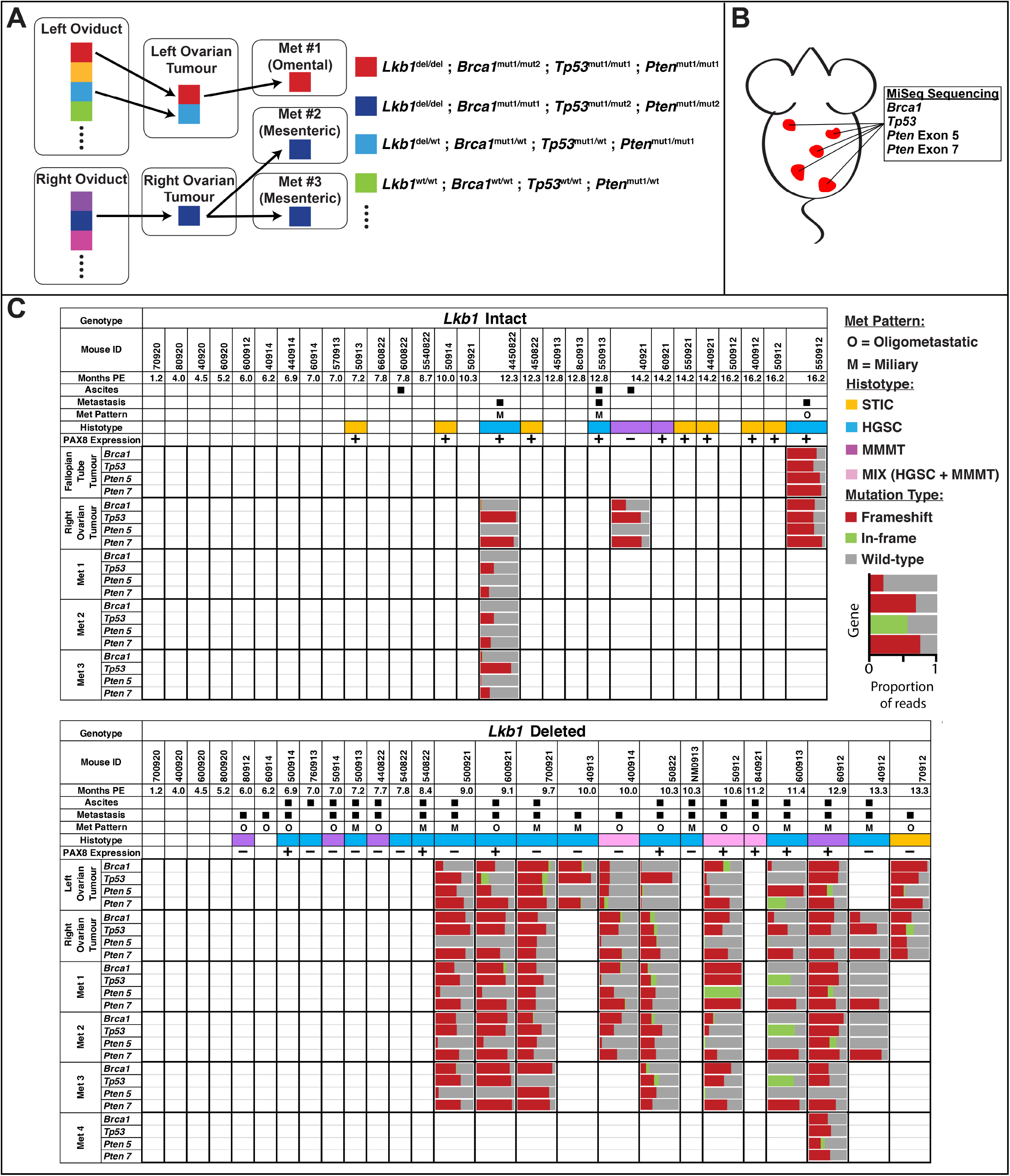
Frame-shift mutations in tumour suppressor genes were predominantly selected during HGSC disease progression. (A) A schematic diagram of clonal development in electroporated cells during ovarian tumour formation and peritoneal metastasis. (B) Tumour samples were isolated from each mouse and analyzed using MiSeq amplicon sequencing of *Brca1, Tp53, Pten* exon 5 and *Pten* exon 7. (C) Overview of phenotypes and mutation patterns in *Lkb1* intact and *Lkb1* deletion mice. Frameshift mutations (red) were predominantly identified in *Brca1, Tp53, Pten* exon 5 and *Pten* exon 7.

Analyses of mutation types in our targeted genes were performed using MiSeq amplicon sequencing of the targeted loci (*Brca1, Tp53, Pten* exon 5 and *Pten* exon 7) in ovarian tumours and 1-3 metastatic lesions from individual mice in *Lkb1* intact and *Lkb1* deletion cohorts (Fig 5B). From the samples analyzed, the majority of the mutations detected in *Brca1, Tp53* and *Pten* were frameshift mutations in both *Lkb1* intact and *Lkb1* deletion mice (Fig 5C). Wild-type reads were also detected, suggesting contamination of normal surrounding cells like stromal and hematopoietic cells, as well as non-mutated alleles in tumour cells. Interestingly, in one *Lkb1* intact mouse (4450822), no *Brca1* mutations were detected in the ovarian tumour and peritoneal metastasis, suggesting that *Brca1* mutations were not absolutely required for tumorigenesis in our model and that a combination of *Tp53* and *Pten* mutations were sufficient to induce HGSC (Fig 5C). On the other hand, some peritoneal metastases in one *Lkb1* deletion mouse (40912) had no mutations in *Brca1* and *Tp53*, suggesting that a combination of loss-of-*Lkb1* and *Pten* mutations would be sufficient to induce HGSC progression in the *Lkb1* deletion cohort (Fig 5C).

### PAX8-papillary tumors developed in the *Lkb1* deletion cohort originated from the OSE and hilum region

Although the *Lkb1* deletion cohort developed HGSC with peritoneal metastasis at a shorter latency and higher penetrance, many tumours in these mice were PAX8- (Fig 4C). Recently, Zhang *et al*. has suggested that both the FTE and OSE in mice have the potential to develop HGSC^5^. Interestingly, in their model, the tumors developed from the FTE were PAX8+ while the tumours derived from the OSE were PAX8-. Therefore, we hypothesized that the PAX8-tumors developed in the *Lkb1* deletion cohort originated from the OSE cells. We carefully examined the distribution of RFP+ cells 5 days PE and found a small number of non-FTE RFP+ cells in adjacent tissues such as the OSE and ovarian hilum, however, the vast majority of electroporated cells were within the distal FTE (Supp Fig.1).

To determine the origin of the tumors formed in the *Lkb1* deletion cohort, we performed salpingectomies (fallopian tube removal) and ovariectomies (ovary removal) immediately after electroporation in 4-6 week old *Lkb1*^flox/flox^; *Rosa-LSLtdTomato* females. Interestingly, the salpingectomized mice still developed large ovarian tumours with widespread peritoneal metastasis at a similar latency to non-salpingectomized mice at 6-7m PE (Table 2). Papillary tumours were found in the OSE and hilum regions 6m PE (Fig 6A). These tumours, along with the ovarian and peritoneal tumours that formed, were PAX8-, WT1+ and CK8+ (Fig 6A).

**Figure 6:**
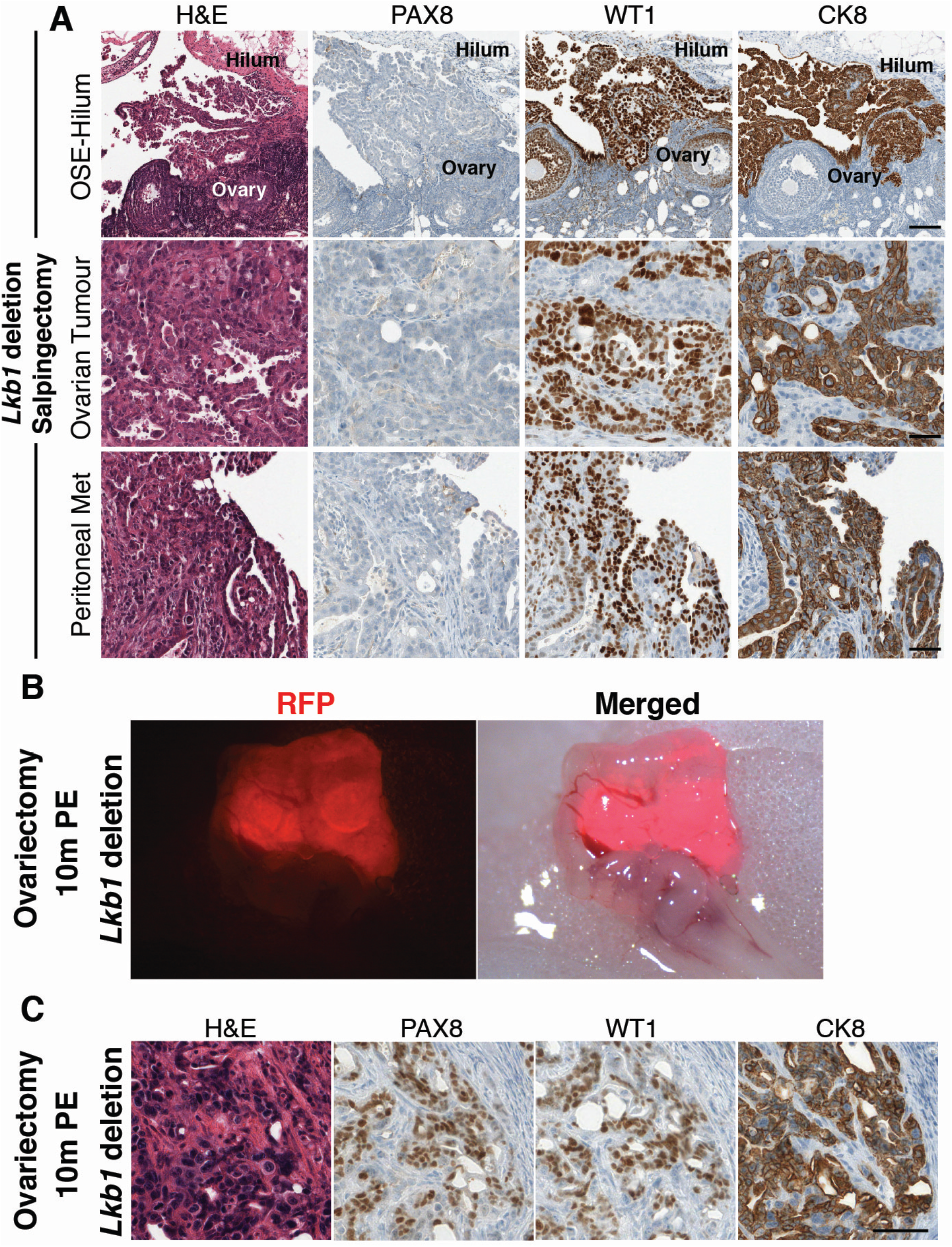
HGSC tumours developed in *Lkb1* deletion mice originated from the OSE and hilum region. (A) H&E, PAX8, WT1 and CK8 immunohistochemical staining in salpingectomized *Lkb1* deletion mouse (top panels). Papillary tumour formation from the OSE-hilum region. These tumours were PAX8-, WT1+ and CK8+. Scale bars 100um. (middle and bottom panels) Ovarian and peritoneal tumours. These tumours were also PAX8-, WT1+ and CK8+. Scale bars 50 μm. (B) RFP+growth within the distal FT of ovariectomized *Lkb1* deletion mouse 10m PE. (C) STIC formed in the fallopian tube in ovariectomized *Lkb1* deletion mouse.

**Table 2.** Summary of disease progression in *Lkb1* deletion mice that have undergone salpingectomy or ovariectomy.

|  |  |  |  |
| --- | --- | --- | --- |
| Month(s) Post Electroporation | Ovarian Tumour Incidence (Per ovary) | Peritoneal Metastasis (Per mouse) | Ascites Formation (Per mouse) |
| **Salpingectomy** |  |  |  |
| 6-8 | 6/10 (60%) | 5/5 (100%) | 5/5 (100%) |
| 9-10 | 4/4 (100%) | 2/2 (100%) | 1/2 (50%) |
| **Ovariectomy** |  |  |  |
| 6-8 | Not applicable | 0/2 (0%) | 0/2 (0%) |
| 9-10 | Not applicable | 0/6 (0%) | 1/6 (17%) |

On the other hand, in the ovariectomized mice, RFP+ growths in the FTs were observed (Fig 6B) however, peritoneal metastasis did not form 6-10m PE (Table 2). It is unknown if the ovary is required for metastatic tumour formation or 10 months was not long enough to allow development of peritoneal metastasis from the FTE. Interestingly, STIC-like precancerous lesions were observed in the distal FTE of ovariectomized mice (Fig 6C).

### Formation of fallopian tube precancerous lesions in both *Lkb1* intact and deletion cohorts with rapid development of papillary tumours in the OSE of *Lkb1* deletion mice

Interestingly, despite the fact that we performed the same electroporation procedure targeting the distal fallopian tube in both *Lkb1* intact and deletion cohorts, the tumours that developed were from different cell populations; FTE and OSE, respectively. This suggested that the OSE and FTE have distinct susceptibility to the two different mutation combinations. To evaluate this, we examined the early cellular responses in the FTE and OSE against each mutation combination.

By 4m PE, STIC-like precancerous lesions were detected in the fallopian tube in both *Lkb1* intact and *Lkb1* deletion cohorts, marked by PAX8, WT1 and CK8 (Fig 7A). The lesions showed epithelial disorganization (Fig 7A,C,E) and consist of RFP+ cells (Fig 7C,E). These lesions expressed PAX8 and an enrichment of membrane phosho-AKT (Fig 7A,E), indicating phosphorylation and activation of AKT. We also found an in increase in cell size in RFP+ STIC-forming cells compared to RFP- or RFP+ non-STIC forming cells, suggesting an increase in mTOR activation in RFP+ STIC-forming cells (Fig 7C,D). This phenotype was further exacerbated with loss-of-*Lkb1* since cell size in RFP+ STIC-forming cells in *Lkb1* deleted mice were larger than *Lkb1* intact cells (Fig 7D). The morphology of these clones was similar to reported human STICs^37,38^.

**Figure 7:**
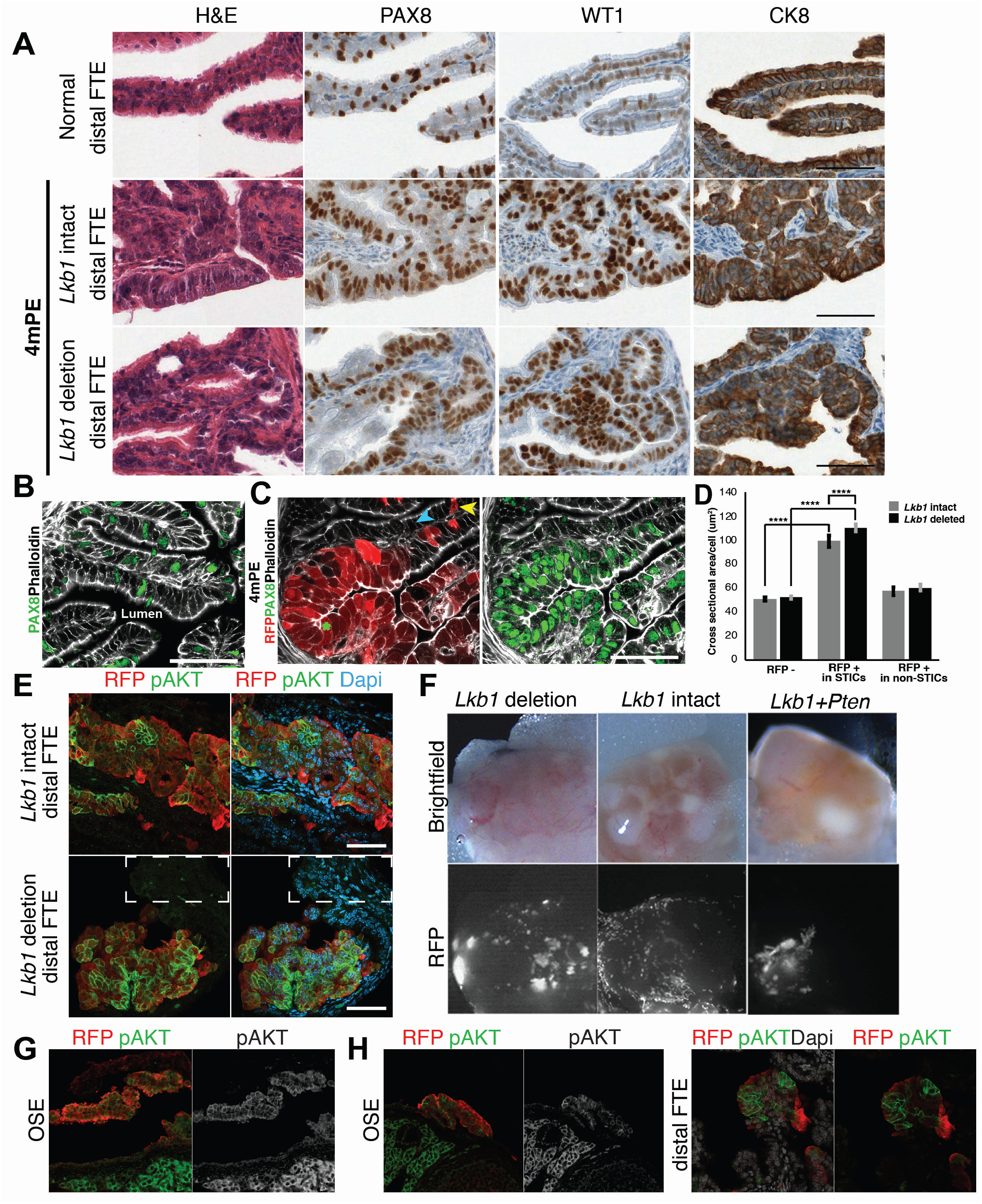
*Lkb1* intact and *Lkb1* deletion FTs developed SITC-like precancerous lesions 4m PE. (A) H&E, PAX8, WT1 and CK8 immunohistochemical staining of normal distal fallopian tube. STIC-like lesions in both *Lkb1* intact and *Lkb1* deletion FTs 4m PE. Scale bars 50 μm. (B) Normal fallopian tube epithelium stained with phalloidin (white) and PAX8 (green). PAX8 secretory cells are randomly distributed throughout the epithelium. Scale bars 50 μm. (C) RFP+ STIC-like lesion within the fallopian tube epithelium stained with phalloidin (white) and PAX8 (green) in *Lkb1* deletion mouse 4m PE. The STIC-like lesions displayed epithelial disorganization and expressed PAX8 (green). Blue arrowhead: RFP-cell, green asterisk: STIC forming RFP+ cell, yellow arrowhead: RFP+ Non-STIC cell. Scale bars 50 μm. (D) A bar-graph of the cross sectional area per cell of RFP-cells, RFP+ cells within STICs and RFP+ cells outside STICs in *Lkb1* intact and *Lkb1* deleted FT STICs. (E) STIC-like lesions in distal FTE of *Lkb1* intact and *Lkb1* deletion mice stained for RFP (red), pAKT (green) and DAPI (blue). RFP+ cells have membrane pAKT. Dotted box outlines RFP-cells that do not show pAKT. Scalebars 50 μm. (F) RFP+ colonies growing on the ovary 2m PE in the *Lkb1* deleted and two gene deleted (*Lkb1* + *Pten*) females but not in the *Lkb1* intact females. (G) Early papillary tumors on the OSE 2m PE in a *Lkb1* deleted female. (H) Two genes deletion (*Lkb1* and *Pten*) was sufficient to form papillary tumours from the OSE and STIC-like lesions in the distal FTE.

To examine the early cellular responses in the OSE, we intentionally targeted the OSE since the number of electroporated cells in the OSE was too small in our standard FTE targeting procedure. As early as 2 months PE, papillary tumour formation from the OSE was observed in the *Lkb1* deletion cohort (Fig 7F,G) while no clonal growth of RFP+ cells was observed in the *Lkb1* intact cohort (Fig 7F). Consistent with a previous study^39^, we found that a combination of loss-of-*Lkb1* and *Pten* alone was sufficient for papillary tumour formation in the OSE (Fig 7F,H). Similar to the STIC-like fallopian tube lesions, RFP+ papillary tumour cells showed an enrichment of membrane pAKT (Fig 7G,H).

These data suggested that the electroporated cells in either the OSE or FTE activated the AKT/mTOR pathway, however, only the cells in the OSE rapidly developed papillary tumours with a high frequency while the cells in the FTE showed only precancerous changes that would need more time for additional oncogenic alterations and progression to develop into HGSC.

## Discussion

In this study, we developed a unique strategy to generate mouse ovarian cancer models using *in vivo* fallopian tube electroporation, CRISPR mediated genome editing and Cre-mediated lineage tracing. Using *in vivo* electroporation, the target area was limited at the distal fallopian tube and ovary, leaving the rest of the female reproductive tract and other tissues/organs unmodified. In contrast to the *Pax8-TetOn-Cre* and *Ovgp1-iCreERT2* mouse models, there was no risk of developing tumours in non-targeted tissues/organs which may necessitate early euthanasia prior to observing symptoms related to HGSC^5,8,34^. The targeting area in this method can also be easily changed by using different sized tweezer type electrodes. The density of electroporated cells can also be modified by changing electroporation parameters and plasmid concentrations. In contrast to systemic Cre activation, which uniformly impacts all Cre expressing cells in targeted tissues, electroporation can create a random mosaic pattern of normal and mutated cells within an epithelium (Fig 1). This mosaic pattern more closely resembles human carcinogenesis which likely occurs in a local sporadic manner. On the other hand, one limitation of the current method is the inability to target specific cell lineages. This can be overcome by controlling Cas9 and Cre expression using a specific cell promoter such as *Ovgp1* or *Pax8*. Various regions of the female reproductive system can be easily targeted with different mutation combinations. This high level of flexibility in choosing gene targets and areas will be useful in developing somatic GEMMs for HGSC^19,21^, which is genetically highly heterogeneous, as well as other types of OCs in which the cell-of-origin and causative mutations are not fully defined^40,41^.

This approach also easily incorporates the Cre reporter system into mouse cancer models to track genetically modified cells. In other HGSC mouse models^3,8,34,39^, peritoneal metastatic tumours are detected when the tumour has grown to a size that can be visible. We identified microscopic metastases in the peritoneum via Cre-mediated RFP expression in cancer cells. Various useful Cre reporter lines are available, such as Confetti^42,43^ and Fucci^44,45^, to track clonal patterns of Cre activated cells and to visualize cell cycles, respectively. These mice will be highly useful to study clonal evolution and mechanisms of peritoneal metastasis formation. Although immunogenicity of transgenes would be a concern from using transgene-based lineage tracing in cancer models, this would be conquered by expressing non-functional transgenes to induce immune tolerance^46^.

The peritoneum is the primary metastatic site for HGSC. Most HGSC patients are diagnosed at advanced stages when tumours cells have already spread to the peritoneal cavity^6,47–49^. Two types of peritoneal metastatic patterns are known; oligometastatic and miliary. It is also known that there is an association between dissemination pattern in the peritoneal cavity and median overall survival in which patients with an upper abdominal/miliary dissemination phenotype have a shorter median overall survival^50,51^. The exact cause for different dissemination patterns in HGSC is unknown but could be driven by a combination of different genomic aberrations and host-tumour interactions. To our knowledge, this is the first mouse model to represent the miliary metastatic pattern of ovarian cancer, which will be highly useful to investigate the complex biology of peritoneal metastasis of HGSC *in vivo*.

Peritoneal metastasis formation is a complex process that involves several necessary steps^52^. First, cancer cells need to detach from the primary site and undergo anoikis resistance in the peritoneal fluid. Secondly, they must land on the peritoneum and integrate into the epithelial lining. Thirdly, the landed micrometastases penetrate a basal membrane of mesothelium to move into the parenchyma. Mesothelial cells are the primary defence against peritoneal metastasis formation in HGSC. They are thin plate-like cells that line the peritoneum and produce a non-adhesive surface on the peritoneal surface through the secretion of glycosaminoglycans and lubricants which allows relatively frictionless movement of the visceral organs^53^. The mechanism for how floating cancer cells, whether they are individual cells or multicellular aggregates, implant onto the peritoneum to form peritoneal metastasis is still not well understood. The attachment and integration of individual cells and multicellular aggregates of human ovarian cancer cell lines were tested using *in vitro* explants of the mouse parietal peritoneum (the peritoneum on the abdominal muscle wall)^54^. An *in vitro* assay recently developed is the mesothelial clearance assay in which ovarian cancer cells are plated on a sheet of immortalized mesothelial cells and integrate into the sheet using acto-myosin dependent cellular contractility^55^. Our observation has confirmed the replacement of mesothelial cells by cancer cells and placing them on the same mesothelial ECM surface, rather than cancer cells directly invading into the organ parenchyma. Some of the landed cancer cells were able to spread and form papillary tumors on the surface peritoneum while others appeared to be penetrating into the organ parenchyma without apparent epithelial mesenchymal transition (EMT). The basal ECM membrane underneath cancer cell clusters were remodeled by the cooperation of the cancer cells and recruited host macrophages. Our observation is consistent with a recent study in which tissue-resident macrophages promote metastatic spread of ovarian cancer^56^. Further studies will be essential to understand the cellular and molecular mechanisms of peritoneal metastasis formation in HGSC.

Cellular pliancy is an emerging concept explaining cell-type specific unique susceptibility against specific oncogenic insults^57^. Similar to cellular competence in reprogramming, preexisting epigenetic states and gene expression patterns of individual cells, which are tightly associated with their lineage and microenvironment, dictate cellular pliancy. Interestingly, in our study, only our *Lkb1* deletion cohort, in which there is a loss-of-*Lkb1* with *Brca1, Tp53* and *Pten* mutations, induced OSE-derived highly metastatic HGSC with a short latency and high penetrance. This indicates that the OSE and FTE have distinct cellular pliancies, although they are anatomically connected and share the same developmental origin; surface coelomic mesodermal cells^58^. The importance of understanding the cell-of-origin of various cancers has been recognized^59^ since the cell-of-origin is able to dictate tumour proliferation, histotype spectrum and subsequently, the immune microenvironment^60^. Our results support this notion and that somatic ovarian cancer GEMMs will be useful to study cellular pliancy in the female reproductive system.

Taken together, our results indicate that different cell types have distinct susceptibilities based on the combination of genetic alterations introduced. Understanding how the cell-of-origin of HGSC contributes to phenotypic heterogeneity, pathophysiology and subsequently, histotype, will be critical in determining correct diagnostic classification and development of effective strategies for prevention, prognosis and treatment of HGSC. Further studies that can elucidate the origin and early pathogenesis for HGSC will be essential for developing novel methods for early detection, treatment and, optimally, prevention of HGSC^9^.

## Materials and Methods

### Mouse lines

All mice were maintained in the Comparative Medicine and Animal Resources Centre at McGill University. All the animal experiments were performed in accordance with institutional guidelines and were approved by the Facility Animal Care Committee (AUP #7843). Conditional *Lkb1*^flox/flox^ mice ^61^ were obtained from the National Cancer Institute (Frederick, MD) and were maintained on a C57BL/6 background. Gt(ROSA)26Sor^tm14(CAG-tdTomato)Hze^/J ^62^ mice were obtained from The Jackson Laboratory (Bar Harbor, ME) and crossed with *Lkb1*^flox/flox^ mice to generate *Lkb1*^flox/flox^; *Rosa-LSLtdTomato* mice.

### *In vivo* oviductal electroporation

All the animal experiments were performed in accordance with institutional guidelines and were approved by the Facility Animal Care Committee (AUP #7843). The surgical procedure was adopted and modified from Takahashi *et al*. ^63^. A dorsal midline skin incision (1cm) was performed and the female reproductive tract (oviduct/ovary) was exposed. Injection procedure was performed under a dissecting scope. Injection solution (approx. 1 μL) contained various plasmids (100-400 ng/ul each) and 0.05% of trypan blue, which was used as a marker for successful injection, was injected into the oviductal lumen using an air-pressure syringe system attached to a micromanipulator. Target region was covered with a small piece of PBS-soaked kimwipe and electroporated with 3mm tweezer type electrodes (BTX Item #45-0487) connected to the BTX 830 or BEX Cuy21 edit2 electroporator. Parameters: 30V, 3 pulses, 1s interval, P. length=50ms, unipolar. Following electroporation, the oviduct/ovary were placed back into their original position and the incisions were sutured.

### Plasmids

PCS2 CreNLS plasmid ^64^ and PX330 plasmid (Addgene # 42230) were used in this study. CRISPR guide sequences (two guides/gene) were designed using ChopChop, CRISPR MIT and Sequence Scan. Each guide efficiency was tested in vitro (Takara protocol) and in vivo. *Tp53* (#2, Exon 5) CATCGGAGCAGCGCTCATGG **TGG**, *Tp53* (#3, Exon 5) CGGAGCAGCGCTCATGGTGG **GGG**, *Pten* (#2, Exon 5) TGTGCATATTTATTGCATCG **GGG**, *Pten* (#3, Exon 7) AGCTGGCAGACCACAAACTG **AGG**, *Brca1* (#1, Exon 6) GCGTCGATCATCCAGAGCGT **GGG**, *Brca1* (#2, Exon 6) GCTACCGGAACCGTGTCAGA **AGG**.

### Mouse dissection protocol

Mice were euthanized in accordance with institutional guidelines and were approved by the Facility Animal Care Committee (AUP #7843). If abdominal ascites was present, the fluid was collected with a 5ml syringe. Surgical scissors were used to cut the parietal peritoneum open and contents of the abdominal cavity were exposed. Images of the peritoneal cavity were taken under a fluorescent dissecting scope and RFP+ tumour samples were carefully dissected out. Each tumour sample (~4-8mm in diameter) was collected for histological and molecular analysis.

### Histopathology, immunohistochemistry and immunofluorescence

Mouse tissue was fixed in 4% PFA at 4°C O/N and embedded in paraffin. H&E staining was performed by the GCRC Histology Core Facility using 5-μm sections. Immunofluorescence staining was done using 5-μm sections. Antigen retrieval was performed in Tris/EDTA buffer for 7 min in pressure cooker. Blocking was done with 1% fish skin gelatin for 1-2hrs at room temperature in a humidified chamber. Primary staining was performed at 4°C O/N. Primary antibodies used: PAX8 (ProteinTech 10336-1-AP), Ki67 (Invitrogen, 14-5698-82), WT1 (Abcam, ab89901), CK8 (Abcam ab53280), acetylated tubulin (Sigma, T7451), podocalyxin (R&D, MAB1556), phosho-mTOR (Cell signaling, #5536), phospho-S6K(Cell signaling, #9205), phospho-4EBP1(Cell signaling, #2855) and phospho-AKT(Cell signaling, #4060). Secondary staining was performed at room temperature for 1-2hrs. Secondary antibodies used: AF-488 phalloidin (Lifetech, A12379), AF-635 phalloidin (Lifetech, A34054), anti-Rabbit 488 (Invitrogen, A21206), anti-Rat 488 (Invitrogen, A21208) and anti-Mouse 488 (Invitrogen, A21202). Phalloidin and DAPI staining was performed during secondary staining period.

### Confocal microscopy

Tissue sections and whole-mounts were mounted in ProLong Gold (Invitrogen, P10144) with a coverslip and were imaged using a Zeiss LSM800 microscope. Laser power thresholds were adjusted manually to give optimal fluorescence intensity for GFP, RFP, Far Red and UV for each antibody combination and applied to each image.

### MiSeq genotyping

MiSeq primers were order with CS1 and CS2 tags attached:

CS1 + Forward primer: 5’- ACACTGACGACATGGTTCTACA + specific forward primer −3’ CS2 + Reverse primer: 5’- TACGGTAGCAGAGACTTGGTCT + specific reverse primer −3’ The following primers were used:*Brca1*-FOR 5’-TGCCCCTTTTTGTTTTACAGT-3’, *Brca1*-REV 5’-AGAACACTTGTCCAGCCACTA-3’ (294bp); *Tp53*-FOR 5’-CTGTGCAGTTGTGGGTCAG-3’, *Tp53*-REV 5’-ACAAATTTCCTTCCACCCGG-3 (258bp); *Pten*(Ex 5)-FOR 5’-GTTGCACAGTATCCTTTTGAAGA-3’, *Pten*(Ex 5)-REV 5’-CAGCTTACCTTTTTGTCTCTGG-3’ (248bp); *Pten*(Ex7)-FOR 5’-AAGAAGTCCTTACATGGGTTGG-3’, *Pten*(Ex7)-REV 5’-TGGCTGAGGGAACTCAAAGT-3’ (290bp). Q5 polymerase was used for all PCR reactions. Initial denaturation at 98°C for 30 seconds, denatured at 98°C for 10 seconds, annealed/amplified at 67°C for 30 seconds, repeated 35x, final extension at 72°C for 2 minutes and held at 4°C. 5μL of PCR sample was run on a 2% agarose gel to visualize expected bands. Remaining PCR products were used for MiSeq amplicon sequencing (Illumina, Spike-in MiSeq PE 250bp) at the Génome Québec Innovation Centre at McGill University. All fastQ sequencing files with a minimum of 100 reads were analyzed using Cas-Analyzer ^65^ and 1% of the total number of reads were filtered out as sequencing errors.

---

**Supplementary Figure 1:**
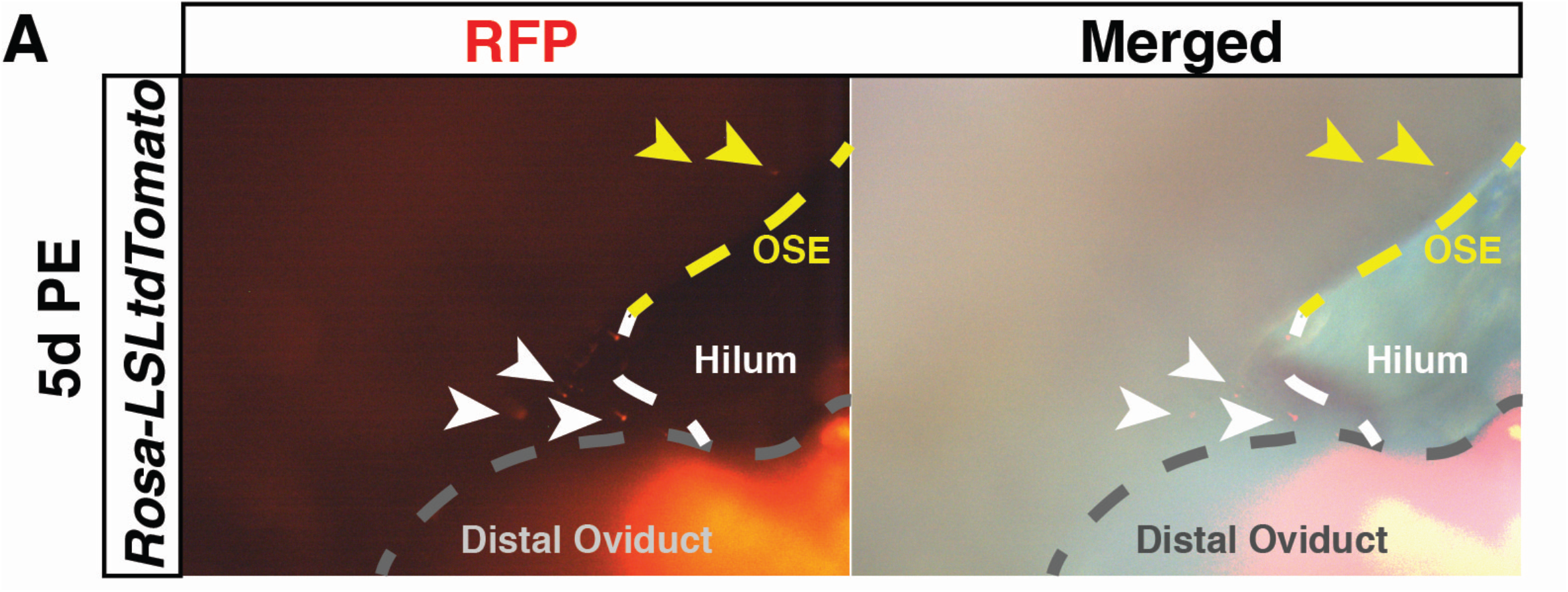
RFP+ electroporated cells in non-fallopian tube epithelial cells. RFP+ electroporated cells on the OSE (yellow arrowheads) and hilum epithelium (white arrowheads) 5 days PE in *Rosa-LSLtdTomato* mouse.

## Author contributions

K.T. executed/analyzed most experiments and wrote/edited the manuscript. M.F., K.H., Y.L. and N.Y. performed some experiments and edited the manuscript. D.F. T-N. TN, D.H. and J.A. provided pathological review. Y.Y. conceived the project and wrote/edited the manuscript

## Acknowledgement

We thank the McGill Goodman Cancer Research Centre Histology and Flow Cytometry Cores. We also thank the McGill Advanced Bioimaging Facility (ABIF). This work was supported by Canadian Cancer Society (CCS) Innovation grant (Haladner Memorial Foundation #704793), CCS i2I grant (# 706320) and Cancer Research Society Operation Grant (#23237). K.T. was supported by MICRTP and Canderel studentships. M.F. was supported by Canderel, CRRD and FRQS postdoc fellowships. K.H. was supported by CRRD and Alexander McFee (Faculty of Medicine) and Gosselin studentships. Y.L was supported by Canderel studentship.

